# Plasmalogen-Independent Neuroprotection: Synthetic alkyl ether lipids activate signaling to mitigate neuroinflammation and enhance cognitive function

**DOI:** 10.64898/2025.12.08.693101

**Authors:** Md Shamim Hossain, Shalman Dipto, Honsho Masanori, Tatsuo Okauchi, Shiro Mawatari, Takehiko Fujino

**Affiliations:** Division of Lipid Cell Biology, Institute of Rheological Functions of Food, Fukuoka, Japan; Department of Neuroinflammation and Brain Fatigue Science, Graduate School of Medical Sciences, Kyushu University, Fukuoka, Japan; Department of applied chemistry, Kyushu Institute of Technology, Fukuoka, Japan

## Abstract

Neuroinflammation is a key driver of neurodegenerative disorders, with chronic activation of the NF-κB pathway contributing to neuronal dysfunction and disease progression. In this study, we report that KIT-8, a synthetic alkyl ether lipid, exerts potent anti-inflammatory and neuroprotective effects despite its inability to be converted into plasmalogens within cells unlike its analog KIT-13, which is known to generate plasmalogens. Remarkably, KIT-8 activates key intracellular signaling pathways, including phosphorylation of ERK and AKT, and enhances brain-derived neurotrophic factor (Bdnf) expression in Neuro2A cells, thereby mimicking the actions of both KIT-13 and natural plasmalogens. Furthermore, KIT-8 significantly improves memory function in mice, underscoring its inherent therapeutic potential. Importantly, the biological activity of KIT-8 suggests that plasmalogen synthesis is not required for its functional effects, introducing a novel concept in ether lipid biology. Supporting this notion, we identified additional alkyl ether lipid analogs, KIT-19 and KIT-20, which also demonstrated anti-inflammatory properties despite lacking the ability to generate plasmalogens. These findings indicate that the ether bond structure alone may be sufficient to confer biological activity. To explore this further, we designed and synthesized a series of structurally distinct alkyl ether lipids (KIT compounds) retaining an ether bond at the sn-2 position. In an LPS-induced inflammatory model using mouse-derived microglial MG6 cells, selected compounds KIT-8, KIT-19, and KIT-20 significantly suppressed NF-κB signaling, reduced p65 nuclear translocation, and downregulated the expression of pro-inflammatory mediators IL-1β and NOS2. Given the cognitive benefits associated with plasmalogens in Alzheimer’s disease, our findings position alkyl ether lipids as promising standalone bioactive molecules, capable of mitigating neuroinflammation and enhancing neuronal function independently of plasmalogen synthesis.

## Introduction

Neuroinflammation is increasingly recognized as a central mechanism in the pathogenesis of various neurodegenerative disorders [1–5]. Activation of inflammatory pathways, particularly the NF-κB cascade leads to the production of cytokines such as interleukin-1β (IL-1β), tumor necrosis factor-alpha (TNF-α), and other pro-inflammatory mediators that contribute to neuronal damage and synaptic dysfunction [6–10]. In Alzheimer’s disease (AD), chronic neuroinflammation is linked to amyloid-β accumulation and tau pathology, which collectively accelerate cognitive decline [1, 11–14]. Similarly, in Parkinson’s disease (PD), sustained inflammatory responses exacerbate dopaminergic neuron loss in the substantia nigra, thereby contributing to motor impairments and non-motor symptoms [15–18]. Moreover, neuroinflammation has been implicated in other age-related conditions, including brain fatigue and cognitive impairment, which are commonly observed in the elderly [19, 20]. These inflammatory processes may not only reduce neuronal viability but also disrupt neuroplasticity and the brain’s ability to recover from injury. Recent studies suggest that the lipid mediators, such as plasmalogens, can mitigate these deleterious effects by exhibiting both neuroprotective and anti-inflammatory properties [3, 11, 21–24]. However, the therapeutic potential of synthetic compounds targeting neuroinflammation can be confounded by their metabolic conversion to plasmalogens and more stable nature, thereby masking their intrinsic biological activities. In this context, a deeper understanding of neuroinflammatory mechanisms across various neurodegenerative and age-related conditions is critical. Such insights can drive the development of targeted therapies that more effectively modulate inflammatory cascades, preserve neuronal function, and ultimately improve clinical outcomes in disorders like AD, PD, and other conditions associated with brain fatigue and cognitive decline.

Plasmalogens, a subclass of alkenyl ether phospholipids, have been shown to exert potent anti-inflammatory and neuroprotective effects [1, 3, 22]. Natural sources of plasmalogens, such as those derived from scallops (sPls), have demonstrated the ability to enhance memory, suppress neuroinflammation, and prevent neuronal apoptosis [3, 13, 21]. Clinical studies further support the cognitive benefits of plasmalogen supplementation in Alzheimer’s disease and mild cognitive impairment [32, 33]. However, their therapeutic application is limited by their metabolic instability and dependence on endogenous enzymatic pathways for conversion and activity. To address these limitations, we have designed and synthesized a series of novel alkyl ether lipids, termed KIT compounds, which retain an ether bond at the sn-2 position but are structurally resistant to conversion into plasmalogens. These synthetic compounds aim to preserve or mimic the biological functions of plasmalogens while offering greater metabolic stability and pharmacological versatility. In this study, we explore the biological potential of these synthetic ether lipids and their therapeutic implications in neuroinflammatory and neurodegenerative contexts.

## Materials and Methods

### Synthesis and Characterization of Alkyl Ether Lipids

A novel series of synthetic alkyl ether lipids (KIT-008, KIT-019, KIT-020) were designed and synthesized. These compounds feature a stable ether linkage at the *sn-2* position of the glycerol backbone but are structurally engineered to resist metabolic conversion into plasmalogens. All reactions were conducted under an inert nitrogen atmosphere and monitored by thin-layer chromatography. Purifications were achieved via silica gel column chromatography. The synthetic route commenced with the stereoselective synthesis of the key alkyl chain precursor, (E)-1-bromooctadec-2-ene. This involved alkylation of a protected propargyl alcohol with 1-bromopentadecane, deprotection to yield octadec-2-yn-1-ol, stereoselective semi-reduction using LiAlH_4_ to give (E)-octadec-2-en-1-ol, and final bromination with PBr_3_. This alkyl bromide was then coupled with the chiral building block (R)-(2,2-dimethyl-1,3-dioxolan-4-yl) methanol under basic conditions. Subsequent acid-catalyzed deprotection yielded the core ether lipid scaffold, (S, E)-3-(octadec-2-en-1-yloxy) propane-1,2-diol. This diol was further elaborated through a sequence of orthogonal protection, phosphorylation, and coupling steps to introduce a phosphoethanolamine head group. Final diversification was accomplished by esterifying the *sn-1* position with distinct polyunsaturated fatty acids docosahexaenoic acid (DHA, for KIT-008), oleic acid (for KIT-019), or arachidonic acid (for KIT-020) using EDC·HCl and DMAP, followed by global deprotection with trifluoroacetic acid. All final compounds were obtained in quantitative yield and their structures were unequivocally confirmed by ^1^H NMR spectroscopy (500 MHz, CDCl_3_), with purity exceeding 95%.

### Cell culture and Reagents

To generate AGPS-knockout (AGPS-KO) HeLa cells, CRISPR/Cas9-mediated gene editing was performed by cloning a guide RNA sequence targeting exon 1 of the human AGPS gene (5’-GTACCAATGAGTGCAAAGCG-3’) into the pX330 vector (Addgene plasmid #42230). The resulting construct was transfected into HeLa cells using Lipofectamine 2000 (Thermo Fisher Scientific, Waltham, MA, USA) according to the manufacturer’s protocol. Following transfection, cells were subjected to single-cell cloning by limiting dilution (0.7 cells/well) in 96-well plates. Successfully established AGPS-KO clones were maintained under standard culture conditions and used for further experiments. AGPS-KO HeLa cells were treated with 20 µg/mL of either KIT-2 or KIT-8 for 48 hours. Total lipid extraction was performed using the Bligh and Dyer method [25], and samples were subjected to further lipid analysis. The Mouse-derived microglial MG6 cells and Neuro2A neuroblastoma cells were cultured in Dulbecco’s Modified Eagle Medium (DMEM; Gibco) supplemented with 10% fetal bovine serum (FBS) and 1% penicillin-streptomycin at 37°C in a humidified incubator with 5% CO_2_. Lipopolysaccharide (LPS) derived from *Escherichia coli* (L2630) was purchased from Sigma-Aldrich and used for stimulation experiments.

### Lipid analysis

Phospholipid classes were separated using high-performance liquid chromatography (HPLC) equipped with an evaporative light scattering detector (ELSD), as previously described [26, 27]. For the specific detection of plasmalogens, acidic hydrolysis was performed using a 2,4-dinitrophenylhydrazine (DNPH)-HCl solution to convert vinyl ether bonds into DNPH-derivatized aldehydes [28]. The resulting DNPH-aldehyde derivatives were separated using an XBridge BEA C18 column (3.0 × 150 mm, 2.5 µm; Waters Corporation) and detected by ultraviolet (UV) absorbance at 356 nm. To quantify plasmalogens containing C18:0 alkenyl chains, liquid chromatography-tandem mass spectrometry (LC-MS/MS) was conducted using a Xevo TQ-S micro mass spectrometer coupled with an ACQUITY UPLC System (Waters), following established protocols [29]. Total lipid extracts were injected into an ACQUITY UPLC BEH C18 column and analyzed using electrospray ionization tandem mass spectrometry (ESI-MS/MS). Data acquisition and quantification were carried out using TargetLynx software (Waters Corporation).

### Immunoblotting

Protein samples were separated by SDS-PAGE and electrotransferred to a polyvinylidene fluoride membrane membrane (Bio-Rad). After blocking in PBS containing 5% non-fat dry milk and 0.1% Tween 20, blots were subjected to immunoblotting with the following antibodies: rabbit polyclonal antibodies to AGPS (Honsho et al. Isolation and characterization of mutant animal cell line defective in alkyl-dihydroxyacetonephosphate synthase. Localization and transport of plasmalogens to post-Golgi compartments. Biochim. Biophys. Acta, 2008, 1783, 1857-1865. PMID: 18571506) and mouse monoclonal antibody to alpha-tubulin (Thermo Fisher Scientific). The p-ERK, p-Akt, and β-Actin were purchased from Cell Signaling Technologies. The immunoblotting with these three antibodies were described in the previous papers [12, 21]. After probing with appropriate HRP-conjugated secondary antibodies, immunoblots were developed with ECL Western blotting detection reagents (GE Healthcare), and scanned with an LAS-4000 Mini luminescent image analyzer (Fujifilm).

### PCR studies

For RT-PCR, total RNA was extracted using TRIzol reagent (Tri-reagent), and 500 ng of RNA was reverse-transcribed using PrimeScript™ RT-PCR Kit (Takara). PCR amplification was carried out using gene-specific primers under standard thermal cycling conditions. PCR products were separated on 2% agarose gels and visualized under UV light. The following mouse primers were used: Bdnf (Forward: ATGGGTGTGAACCACGAGAA, Reverse: AGTTGTCATGGATGACCTTGG), IL-1β (Forward: *AAAGCTCTCCACCTCAATGG*, Reverse: *AGGCCACAGGTATTTTGTCG*), Gapdh (Forward: ACTCACCTCTTCAGAACGAATTG, Reverse: CCATCTTTGGAAGGTTCAGGTTG), and β-Actin (Forward: CACTGTGCCCATCTACGA, Reverse: CAGGATTCCATACCCAAG).

### ELISA assays to detect secreted IL-1β in microglial cells

MG6 microglial cells were seeded at a density of 5,000 cells/well in 100 µL of DMEM in 96-well plates. After 24 hours of incubation, cells were pretreated with KIT-13 (5 µg/mL), KIT-19 (5 µg/mL), or KIT-20 (5 µg/mL) for 12 hours. Subsequently, lipopolysaccharide (LPS) was added at a concentration of 1 µg/mL for 6 hours to induce an inflammatory response. Following treatment, culture supernatants were collected for cytokine quantification by ELISA. ELISA was performed following a standard sandwich protocol provided by the supplier (Mouse IL-1 beta DuoSet ELISA, R&D, DY401). Briefly, 96-well plates were coated with 100 µL of capture antibody diluted in PBS and incubated overnight at room temperature. Plates were washed three times with Wash Buffer (400 µL/well) and blocked with 300 µL of Reagent Diluent for 1 hour at room temperature. After washing, 100 µL of each sample (20 µl of cell supernatant protein) or standard was added per well and incubated for 2 hours at room temperature. Plates were washed, and 100 µL of detection antibody diluted in Reagent Diluent was added and incubated for 2 hours. After another washing step, 100 µL of Streptavidin-HRP was added and incubated for 20 minutes in the dark. Plates were washed again, and 100 µL of Substrate Solution was added for 20 minutes at room temperature. The reaction was stopped with 50 µL of Stop Solution, and absorbance was measured at 450 nm with wavelength correction at 540 nm using a microplate reader.

### Morris Water Maze Test

The experimental design and all animal procedures were approved by the Ethical Committee for Animal Use at the Institute of Rheology Function of Foods, as described in our previous study [30]. The Morris Water Maze (MWM) and subsequent probe trials were conducted in accordance with established protocols [13, 23, 30], with minor modifications to enhance consistency and reproducibility. Male C57BL/6J mice (9 weeks old, n = 10 per group) were housed under standard laboratory conditions with a 12-hour light/dark cycle and ad libitum access to food and water unless otherwise specified. The water maze consisted of a circular white pool (120 cm in diameter) filled with water maintained at 23-25°C and rendered opaque using non-toxic white tempera paint. A hidden platform (10 cm in diameter) was placed 1.5 cm below the water surface in one of the quadrants. The experimental groups included a control group (no treatment), an LPS-treated group (intraperitoneal injection of 250 µg/kg once daily for 7 consecutive days), and LPS + KIT-treated groups, which received the same LPS regimen. For KIT pre-treatment, mice were orally administered either KIT-8 or KIT-20 at a dose of 10 mg/50 kg once daily for 30 days prior to the initiation of LPS injections. Behavioral training (learning phase) was conducted over six consecutive days with six trials per day. In each trial, mice were released from randomized starting positions and allowed 60 seconds to locate the submerged platform. Mice that failed to find the platform within the allotted time were gently guided to it and allowed to remain for 15 seconds. Twenty-four hours after the final training session, a probe test was performed to evaluate spatial memory retention. The platform was removed, and mice were allowed to swim freely for 60 seconds. Latency to reach the previous platform location and the time spent in the target quadrant were recorded and analyzed using automated tracking software. Both the average latency during the learning phase and performance in the probe test were used as key indicators of spatial learning and memory function.

### Immunohistochemistry

Immunohistochemistry (IHC) was performed according to previously described methods with slight modifications [2, 13]. Briefly, mice were deeply anesthetized and transcardially perfused with phosphate-buffered saline (PBS), followed by 4% paraformaldehyde (PFA) in PBS. Brains were carefully removed, post-fixed in 4% PFA overnight at 4°C, and then cryoprotected in 30% sucrose solution until they sank. Coronal brain sections (10 µm thick) were prepared using the cryostat at −18°C and stored in cryoprotectant solution at −70°C until staining. Free-floating sections were washed in PBS, then blocked with 5% normal goat serum containing 0.2% Triton X-100 in PBS for 1 hour at room temperature to reduce nonspecific binding. Sections were incubated overnight at 4°C with the following primary antibodies: rabbit anti-Iba1 (1:1000, Wako), Cy3-conjugated anti-GFAP (1:4000, Millipore), mouse anti-NeuN (1:1000, Millipore), and rabbit anti-iNOS (1:500, Cell Signaling Technology). For unconjugated primary antibodies, sections were incubated the next day with appropriate Alexa Fluor-conjugated secondary antibodies (1:2000; Invitrogen) for 2 hours at room temperature. After final washes in PBS, sections were mounted onto glass slides with Vectashield mounting medium containing DAPI (Vector Laboratories) for nuclear counterstaining. Images were captured using a Zeiss Axioskop 2 fluorescence microscope equipped with appropriate filter sets and a high-resolution digital camera. Image analysis and quantification were performed using ImageJ software (NIH).

## Results

### KIT Compounds Differentially Affect Plasmalogen Synthesis in AGPS-KO HeLa Cells

We first established the impact of KIT compounds on plasmalogen biosynthesis using AGPS-KO HeLa cells. As shown in Figure 1A, Western blot analysis confirmed the absence of AGPS protein in these cells, validating the knockout model. Treatment with KIT2, but not KIT8, led to a significant restoration of plasmalogen levels. Quantitative analysis using HPLC-ELSD revealed that cells incubated with KIT2 exhibited markedly elevated plasmalogen (Pls, C18:0) levels compared to those treated with KIT8 (Figure 1B). This observation was further supported by the detection of plasmalogen-derived aldehydes in KIT2-treated cells using acidic hydrolysis with DNPH-HCl followed by HPLC-UV analysis at 356 nm (Figure 1C). Moreover, LC-MS/MS analysis confirmed the presence of plasmalogens containing C18:0-alkenyl chains exclusively in cells treated with KIT2 (Figure 1D). Quantification of plasmalogen levels (Figure 1E) substantiated that while KIT2 effectively reinstated plasmalogen synthesis, KIT8 did not, underscoring a functional divergence between these compounds with respect to plasmalogen formation.

**Figure 1.**
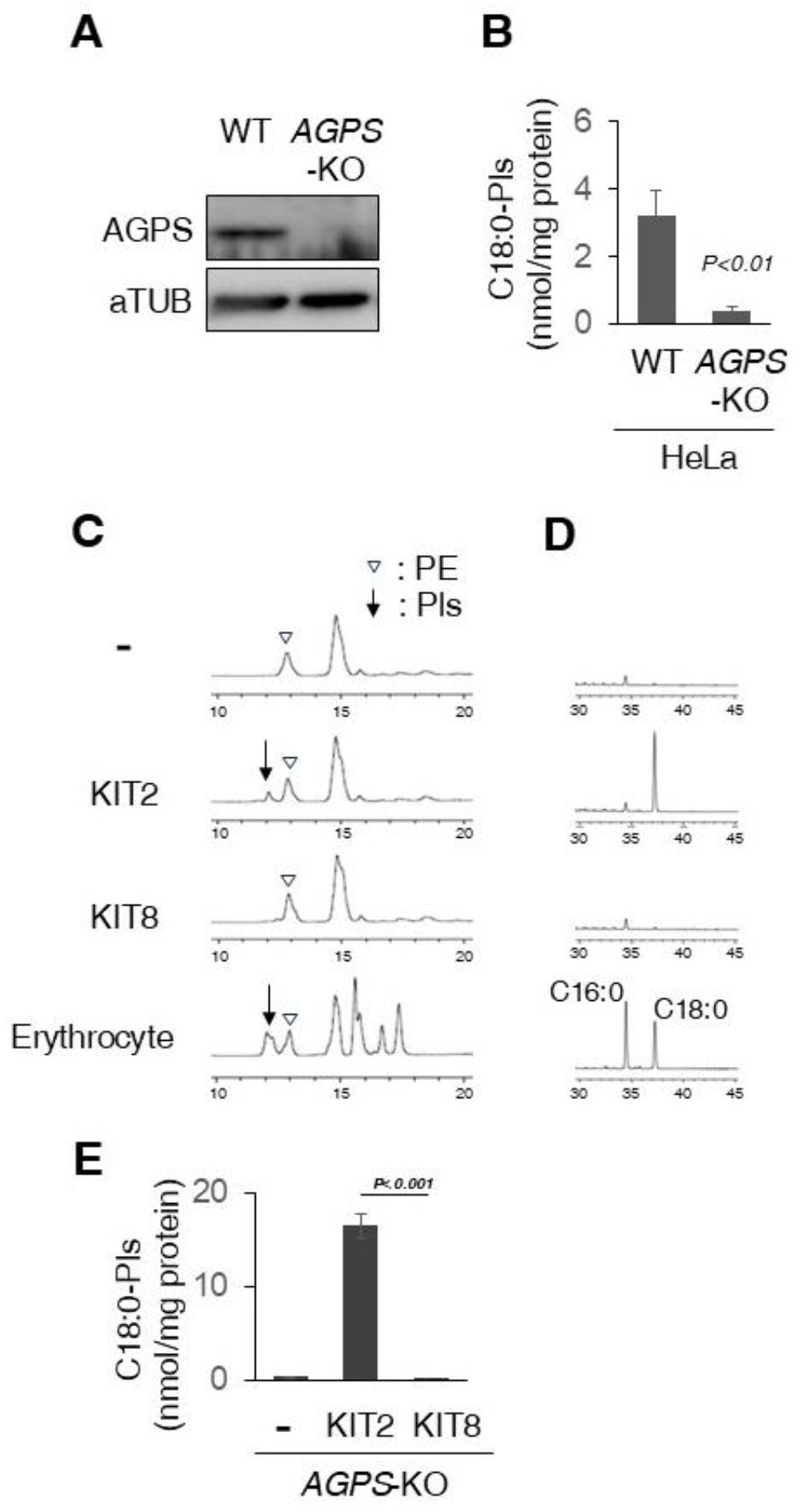
Plasmalogen synthesis in AGPS-KO HeLa cells in response to KIT2 and KIT8 treatment. **(A)** Western blot analysis showing the absence of AGPS expression in AGPS-KO cells. The data represents three independent experiments (n=3). **(B)** Plasmalogen (Pls) levels (C18:0) were quantified using HPLC-ELSD after AGPS-KO cells were cultured with KIT2 or KIT8. **(C)** Aldehydes derived from plasmalogens were found in KIT-2 treated cells when detected by acidic hydrolysis using DNPH-HCl and analyzed by HPLC-UV at 356 nm. PE stands for phosphatidylethanolamine. **(D)** LC-MS/MS analysis confirming plasmalogens containing C18:0-alkenyl chains in AGPS-KO cells treated with KIT2 but not KIT8. **(E)** Quantification of plasmalogen levels in KIT2 and KIT8 treated AGPS KO Hela cells. Data are represented as mean ± SD (n = 3). Bars represent the mean, and error bars indicate the standard error of the mean (SEM). P values were calculated using Student’s t-test.

### Activation of GPR21 Signaling and Upregulation of Bdnf Expression in Neuro2A Cells

To elucidate the intracellular signaling pathways activated by KIT compounds, Neuro2A cells were treated with KIT-8, KIT-13, or scallop-derived plasmalogens (sPls) at a concentration of 5 µg/mL for 2 hours. Western blot analysis revealed a significant increase in the glycosylated form of GPR21, along with enhanced phosphorylation of ERK and AKT, compared to untreated controls (Figure 2A). These results align with our previous findings showing that sPls promote glycosylation of GPR21 [12], a G protein-coupled receptor proposed as a potential plasmalogen sensor [31]. To evaluate whether KIT-8 could mimic this effect, we examined GPR21 glycosylation as a functional marker of receptor activation. Densitometric analysis confirmed that the observed changes were statistically significant (Figure 2B). Additionally, RT-PCR analysis demonstrated that a 6-hour treatment with KIT-8, KIT-13, or sPls resulted in a marked upregulation of Bdnf mRNA levels (Figure 2C), with quantitative analysis indicating that KIT-8 stimulated Bdnf expression to a degree comparable to that of KIT-13 and sPls (Figure 2D). These findings suggest that KIT-8, despite lacking the ability to generate plasmalogens, can effectively activate GPR21-mediated signaling pathways and promote neurotrophic gene expression.

**Figure 2.**
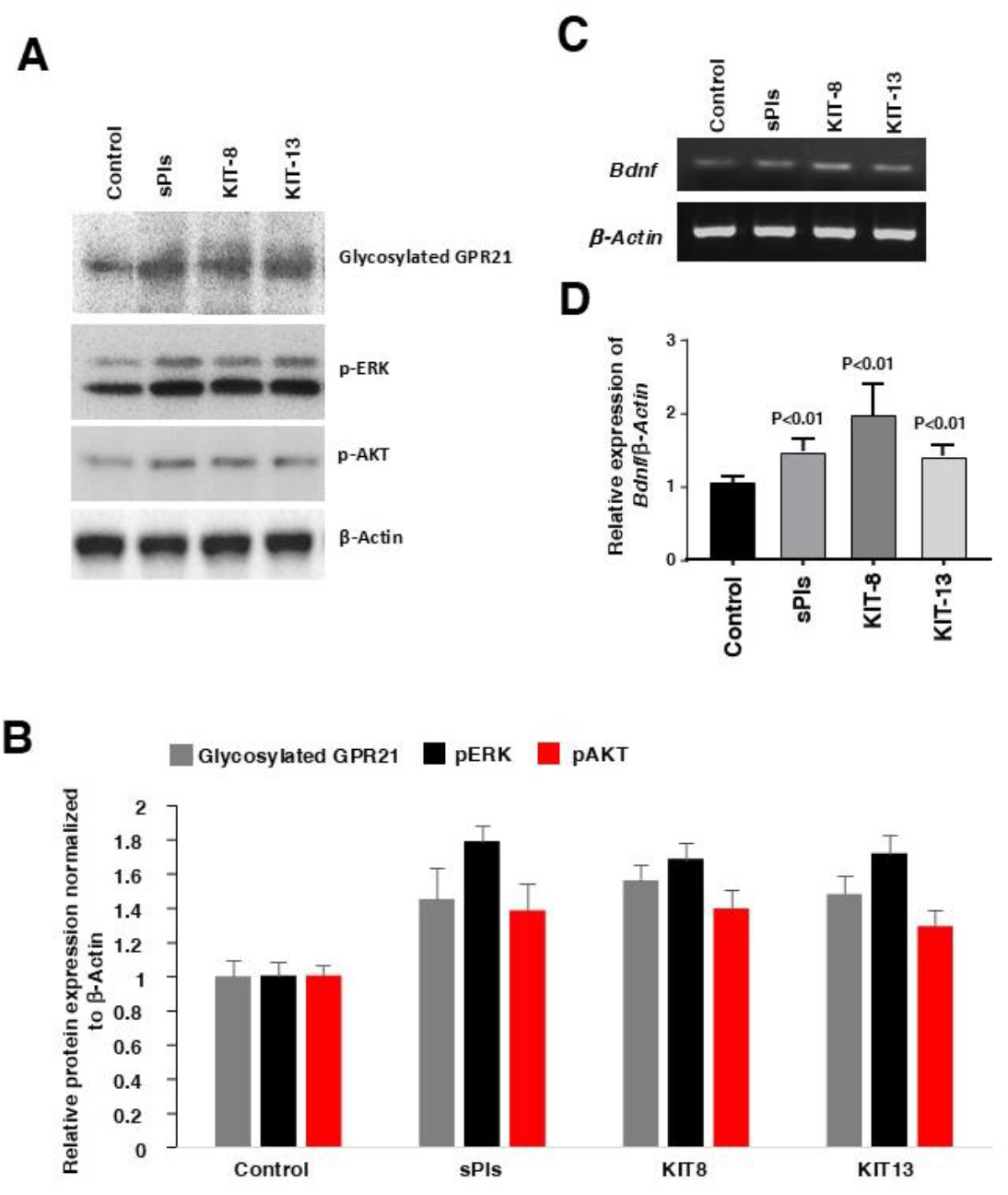
KIT Compounds and Scallop-Derived Plasmalogens Activate GPR21 Signaling and Induce Bdnf Expression in Neuro2A Cells. **(A)** Western blot analysis of glycosylated GPR21, phosphorylated ERK (p-ERK), and phosphorylated AKT (p-AKT) in Neuro2A cells treated with 5 µg/ml of KIT8, KIT13, or scallop-derived plasmalogen (sPls) for 2 hours. Representative blot images show increased levels of glycosylated GPR21, p-ERK, and p-AKT compared to untreated control. β-Actin was used as a loading control. The data represent three independent experiments. **(B)** Densitometric quantification of Western blot results (Image J software) from panel A. All three treatments significantly upregulated glycosylated GPR21, p-ERK, and p-AKT levels (n=3). **(C)** RT-PCR analysis of Bdnf mRNA expression in Neuro2A cells treated with 5 µg/ml of KIT8, KIT13, or sPls for 6 hours in DMEM containing 2% FBS. The upper gel image shows Bdnf expression, while the lower panel shows β-Actin as a loading control. **(D)** Quantitative analysis of Bdnf mRNA expression based on RT-PCR results in panel C. KIT8 significantly increases Bdnf expression, with levels comparable to those induced by KIT13 and sPls treatments (n=3). The bars of panel B and D indicate average value of three independent experiments (n=3), and error bars indicate SEM. The P values were calculated using one-way ANOVA followed by Bonferroni post hoc tests.

### Inhibition of LPS-Induced Pro-inflammatory Responses in MG6 Microglial Cells

Plasmalogens and the synthetic analog KIT-13 have previously been shown to exert anti-inflammatory effects in LPS-induced inflammation models using mouse microglial cells. To determine whether other KIT compounds share similar properties, we investigated the anti-inflammatory potential of KIT-8, KIT-19, and KIT-20 in the same LPS-stimulated microglial model. As shown in Figure 3B, pretreatment of MG6 cells with KIT-8 or scallop-derived plasmalogens (sPls) at 5 µg/mL for 12 hours significantly suppressed the LPS-induced upregulation of *Il1b* mRNA, as assessed by RT-PCR. Furthermore, ELISA analysis of culture supernatants revealed that pretreatment with KIT-13, KIT-19, or KIT-20 markedly reduced IL-1β protein secretion following LPS stimulation (Figure 3C). These findings indicate that, like plasmalogens and KIT-13, the structurally distinct alkyl ether lipids KIT-8, KIT-19, and KIT-20 also exert robust anti-inflammatory effects in activated microglia, both at the transcriptional and protein levels.

**Figure 3.**
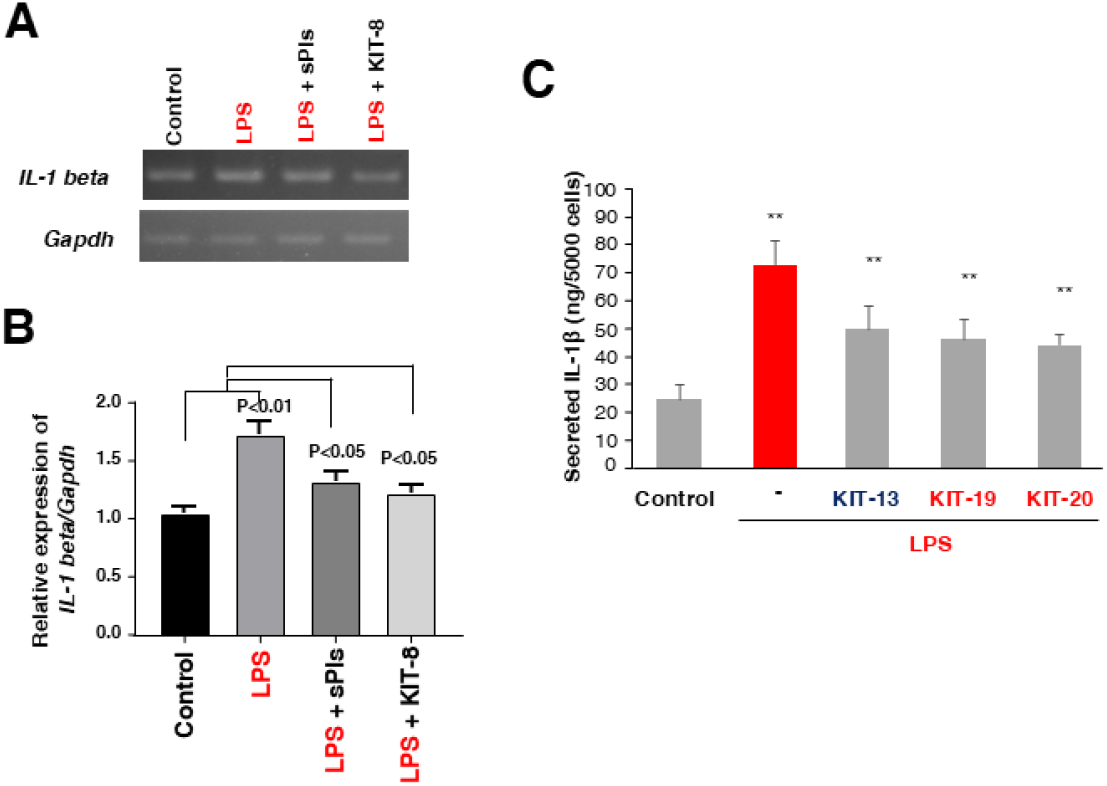
KIT compounds and sPls suppress LPS-induced IL-1β expression and secretion in MG6 microglial cells. **(A-B)** Mouse microglial MG6 cells were pretreated with KIT8 or scallop-derived plasmalogen (sPls) at 5 µg/ml for 12 hours, followed by stimulation with lipopolysaccharide (LPS; *E. coli*, 1µg/ml) for 6 hours. Total RNA was extracted and subjected to RT-PCR analysis to examine the expression of the pro-inflammatory cytokine Il1β (IL-1β). The upper gel panel shows Il1β mRNA expression, and the lower panel shows *Gapdh* as an internal control. The bar graph quantifies Il1β mRNA levels normalized to *Gapdh*. Results demonstrate that pretreatment with KIT8 and sPls attenuates LPS-induced Il1β expression. Data are representative of three independent experiments (n=3). **(C)** ELISA analysis of secreted IL-1β protein levels in the culture supernatants of MG6 cells. Cells were pretreated with KIT13, KIT19, or KIT20 (5µg/ml) for 12 hours before LPS stimulation (1µg/ml, 6 hours). The LPS-induced increase in IL-1β secretion was significantly attenuated by pretreatment with each of the KIT compounds, indicating their anti-inflammatory effects at the protein level. The data shows as average of 5 experiments in each group (n=5) and the error bar indicates SEM. The P values (**, *p*<0.05) including panel B were calculated using one-way ANOVA followed by Bonferroni post hoc tests.

### KIT-8 and KIT-20 Rescue Cognitive Performance and Reduce Neuroinflammatory Stress in the Hippocampus

To evaluate the neuroprotective effects of KIT-8 and KIT-20 in an LPS-induced neuroinflammatory mouse model, we assessed both cognitive function and hippocampal inflammation in the same animals. LPS-induced memory impairment in mice was previously published [23, 30]. Behavioral performance was first evaluated using the Morris water maze test. During the learning phase, mice treated with LPS exhibited significantly longer escape latencies across six days of training compared to the control group, indicating impaired spatial learning (Figure 4A). In contrast, mice co-treated with either KIT-8 or KIT-20 demonstrated improved learning, with significantly reduced escape times approaching those of control animals. Following training, memory retention was assessed using the probe test. LPS-treated mice spent significantly less time in the target quadrant, reflecting deficits in spatial memory (Figure 4B). However, co-administration of KIT-8 or KIT-20 restored performance, as these mice spent significantly more time in the target zone, comparable to the control group. To determine whether these cognitive improvements were associated with reduced neuroinflammation, we conducted immunohistochemical analyses of the hippocampus from the same animals. LPS administration induced robust expression of the inflammatory marker iNOS, particularly in NeuN-positive neurons (Figure 4C, arrowheads), suggesting neuron-associated inflammatory stress. Strikingly, co-treatment with KIT-8 or KIT-20 substantially reduced iNOS expression, preserving neuronal integrity. Quantitative analysis confirmed that iNOS signal intensity was significantly elevated in LPS-treated mice and was markedly attenuated by both KIT-8 and KIT-20 treatments (Figure 4D).

**Figure 4.**
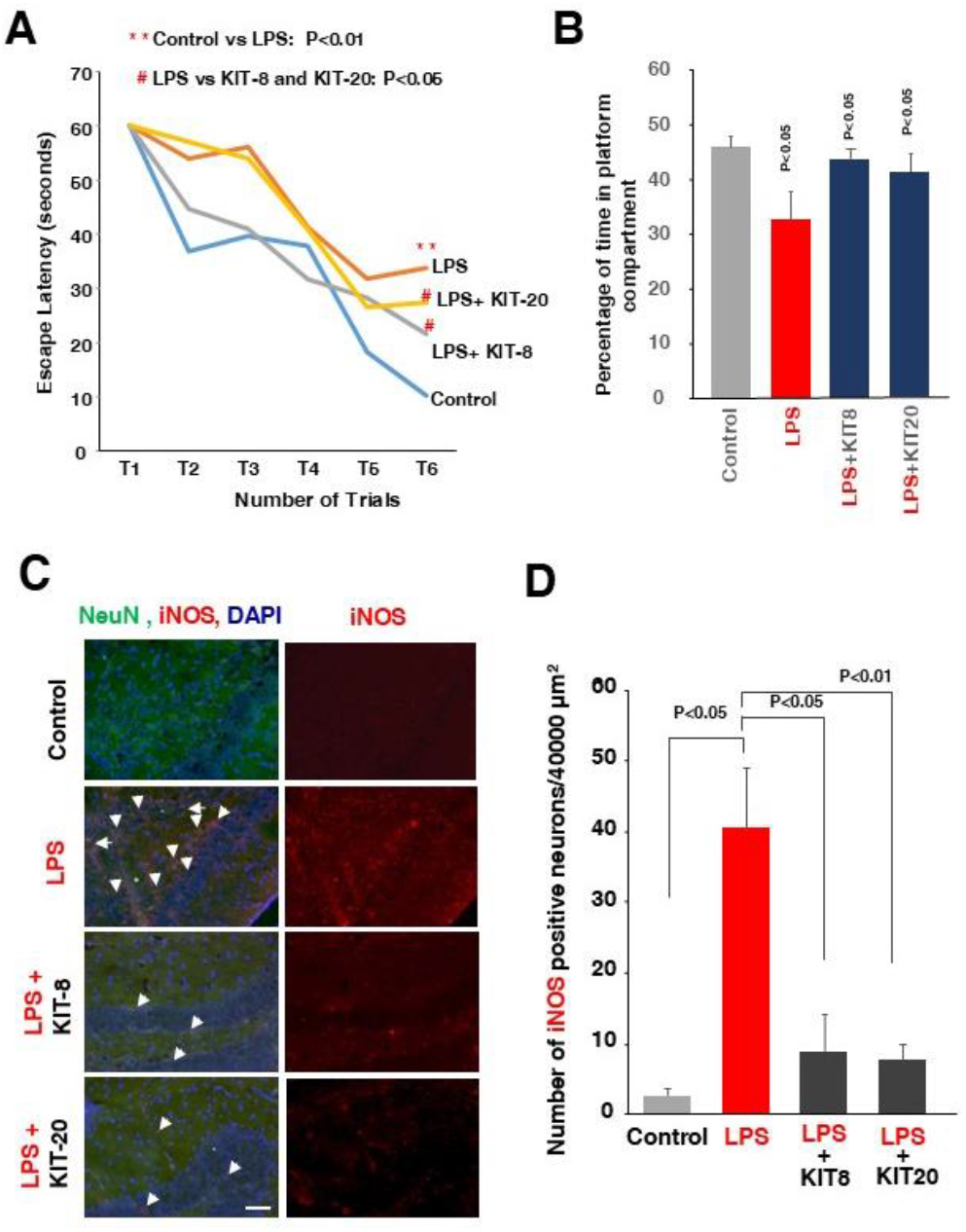
KIT8 and KIT20 attenuate LPS-induced cognitive deficits and neuroinflammation in mice. **(A)** *Morris water maze learning test*. Latency to reach the hidden platform was measured over consecutive 6 days (T1-T6, indicates one trials per day for 6 days) of training. Mice (10 in each group) treated with LPS showed significantly longer escape latency on day 6 compared to controls, indicating impaired learning. The pretreatments with KIT8 and KIT20 markedly improved performance with the same LPS treatments. **(B)** *Morris water maze probe test*. Following the training period, the hidden platform was removed to assess spatial memory retention. Time spent in the target quadrant (where the platform was previously located) was measured. LPS-treated mice spent significantly less time in the target area, suggesting impaired memory retention, whereas KIT8 and KIT20 co-treated mice showed restored performance comparable to controls. **(C)** *Immunohistochemical analysis of brain sections*. Sections were stained for NeuN (neuronal marker, green), iNOS (inflammatory marker, red), and DAPI (nuclei, blue). LPS administration induced strong iNOS expression, particularly in NeuN-positive neurons (white arrowheads). KIT8 and KIT20 co-treatment reduced iNOS expression and preserved neuronal integrity. Scale bar: 50 µm. **(D)** *Quantification of iNOS expression from panel C*. LPS significantly increased iNOS-positive signal intensity in the hippocampus, which was significantly attenuated by KIT8 and KIT-20 co-treatment. Data represent mean ± SEM (n = 3-4 mice per group). *p < 0.05, **p < 0.01 vs. Control; #p < 0.05 vs. LPS. The P values were calculated using one-way ANOVA followed by Bonferroni post hoc tests.

Collectively, these results demonstrate that KIT-8 and KIT-20 not only prevent LPS-induced cognitive decline but also suppress neuroinflammation in the mice brain, highlighting their potential as neuroprotective agents in inflammatory brain disorders like that of the natural plasmalogens (sPls) and plasmalogen producing derivatives KIT-13.

### KIT-8 and KIT-20 Suppress LPS-Induced Astrocyte and Microglial Activation in the Hippocampus

In our previous studies, KIT-13 was shown to attenuate neuroinflammation by reducing astrocytic and microglial activation in the brains of LPS-injected mice [30]. To determine whether structurally related compounds KIT-8 and KIT-20 exhibit similar anti-inflammatory potential, we evaluated glial activation in the hippocampus following LPS administration. Immunofluorescence staining using GFAP (a marker for reactive astrocytes) and Iba1 (a marker for activated microglia) revealed that LPS treatment markedly increased the expression of both markers, indicative of pronounced glial activation (Figure 5A). High-magnification images highlighted typical features of reactive gliosis, including hypertrophied astrocytic processes and elevated microglial density. Strikingly, co-treatment with KIT-8 or KIT-20 significantly reduced the immunoreactivity of both GFAP and Iba1, suggesting a strong inhibitory effect on LPS-induced gliosis. Quantitative analysis of fluorescence intensities further confirmed that the upregulation of both glial markers was significantly attenuated by the KIT compounds (Figure 5B). These results demonstrate that KIT-8 and KIT-20 effectively suppress astrocytic and microglial activation in the hippocampus, like the effects previously observed with KIT-13. Notably, despite their inability to generate plasmalogens like KIT-13, both KIT-8 and KIT-20 exhibited comparable anti-inflammatory efficacy. This supports the concept that the alkyl ether lipid structure itself is sufficient to confer neuroprotective and anti-inflammatory activity, independent of plasmalogen biosynthesis.

**Figure 5.**
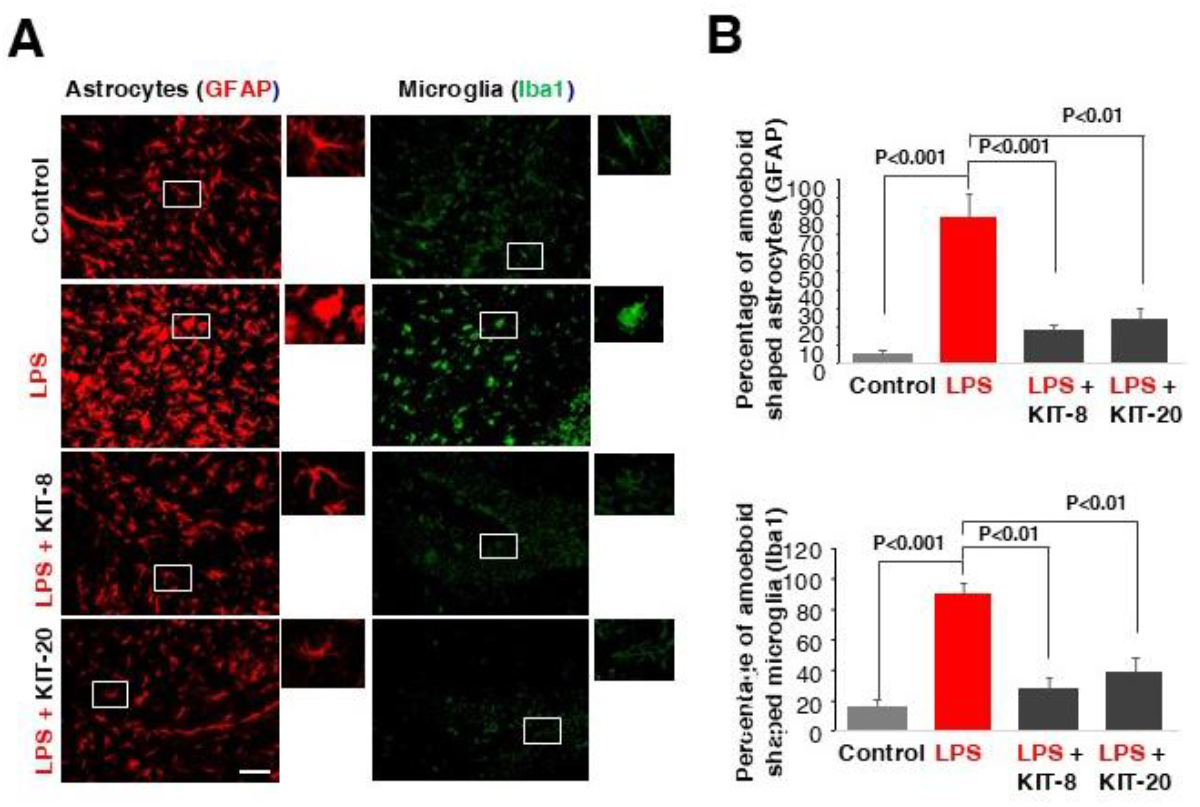
KIT8 and KIT20 attenuate LPS-induced astrocyte and microglial activation in the mouse hippocampus. **(A)** Representative immunofluorescence images of hippocampal sections showing GFAP (red; astrocyte marker) and Iba1 (green; microglial marker) staining in control, LPS-treated, and LPS + KIT8 or LPS + KIT20 co-treated groups. LPS treatment markedly increased GFAP and Iba1 expression, indicating activation of astrocytes and microglia. Co-treatment with KIT8 or KIT20 reduced GFAP and Iba1 immunoreactivity, suggesting an anti-inflammatory effect. Enlarged insets (right panels) highlight morphological changes in glial cells. **(B)** Quantification of GFAP (top graph) and Iba1 (bottom graph) fluorescence intensity. LPS significantly elevated both markers compared to control, while KIT8 and KIT20 significantly attenuated this increase. Data represent mean ± SEM from three independent experiments. **\***p < 0.05, **\*\***p < 0.01 vs. control; #p < 0.05 vs. LPS. Scale bar = 50 µm. The P values were calculated using one-way ANOVA followed by Bonferroni post hoc tests.

## Discussion

Our study reveals a novel paradigm in ether lipid biology by demonstrating that synthetic alkyl ether lipids, such as KIT-8 and KIT-20, possess potent anti-inflammatory and neuroprotective properties independent of their ability to generate plasmalogens. Traditionally, the biological benefits of ether lipids have been largely attributed to their conversion into plasmalogens, the lipids essential for membrane structure and neuronal function. However, our findings challenge this view by showing that the ether bond structure itself, even in the absence of plasmalogen biosynthesis, can mediate significant physiological effects.

KIT-8, a synthetic ether lipid that cannot restore plasmalogen levels, mimicked the biological activity of plasmalogen-producing analogs like KIT-13 in multiple in vitro and in vivo models. In Neuro2A cells, KIT-8 activated intracellular signaling pathways, including ERK and AKT phosphorylation, and significantly upregulated Bdnf mRNA expression, suggesting a role in promoting neuronal survival and plasticity. Importantly, this effect was accompanied by increased glycosylation of GPR21, a G protein-coupled receptor recently implicated as a putative sensor for plasmalogens [3, 12, 31]. The parallel activation of GPR21 and its downstream signaling by KIT-8 points to a receptor-mediated mechanism of action, where ether lipids may act as functional ligands, triggering neuroprotective cascades like those initiated by natural plasmalogens.

In mouse microglial MG6 cells, KIT-8, KIT-19, and KIT-20 each significantly suppressed LPS-induced inflammatory responses. These compounds reduced the expression of key pro-inflammatory markers, including Il1β, and decreased IL-1β protein (pro-inflammatory cytokine) secretion, aligning with the known effects of plasmalogens and KIT-13. Strikingly, the anti-inflammatory activities of KIT-8 and KIT-20 were recapitulated in vivo, where they ameliorated LPS-induced cognitive deficits and reduced neuroinflammatory stress in the hippocampus. Mice treated with these compounds exhibited improved performance in the Morris water maze test and showed markedly lower hippocampal expression of iNOS (pro-inflammatory marker). In addition, reduced inflammatory phenotypes like amoeboid shaped astrocytes (GFAP positive) and microglia (Iba1 positive), indicating that these KITs can attenuate neuroinflammation and glial activation when ingested orally. These data collectively highlight the capacity of non-plasmalogen-producing ether lipids to rescue cognitive function and preserve neuronal integrity under inflammatory conditions. Like KIT-13, oral administration of KIT-8 and KIT-20 exerted anti-inflammatory effects in the mouse brain, raising the important question of whether these lipids can cross the blood–brain barrier (BBB). Although the ability of KIT compounds to directly penetrate the BBB remains uncertain, it is also plausible that their biological activity could be mediated through peripheral mechanisms. For instance, these compounds may activate G protein-coupled receptors (GPCRs) in peripheral tissues. In our previous work, we identified GPR21 expression in natural killer (NK) cells, suggesting that these ether lipids might act on peripheral immune cells, thereby indirectly modulating brain function. Further investigation will be required to determine the precise mechanisms of action and the potential for BBB permeability of these compounds.

These findings shift the therapeutic focus from plasmalogen replenishment to a broader exploration of ether lipid bioactivity. The fact that KIT-8 and structurally related compounds exert their effects without restoring plasmalogen levels implies a plasmalogen-independent mechanism that may involve direct interaction with membrane receptors such as GPR21. This ligand-like activity provides a compelling molecular basis for the anti-inflammatory and neuroprotective actions of ether lipids and expands our understanding of their functional repertoire. The therapeutic implications of these results are significant. While plasmalogen supplementation has shown promise in improving cognitive function in Alzheimer’s disease and MCI [32, 33], our findings reveal that synthetic ether lipids such as KIT-8 and KIT-20 can exert neuroprotective effects independently of plasmalogen biosynthesis. This suggests a broader therapeutic potential for ether lipid-based interventions, especially in conditions where plasmalogen synthesis is impaired. Developing synthetic ether lipids that do not require conversion to plasmalogens could overcome the limitations associated with plasmalogen restoration therapies. Developing synthetic ether lipids that do not require conversion to plasmalogens could overcome the limitations associated with plasmalogen restoration therapies, particularly in disorders like rhizomelic chondrodysplasia punctata or Alzheimer’s disease, where plasmalogen biosynthesis is impaired. Our study suggests that the ether bond structure alone may serve as a bioactive motif, capable of engaging key cellular signaling networks and offering a novel strategy for mitigating neuroinflammation and cognitive decline.

In summary, our work identifies KIT-8 and KIT-20 as functionally active ether lipids that recapitulate the neuroprotective and anti-inflammatory effects of plasmalogens without relying on plasmalogen synthesis. This highlights the therapeutic promise of alkyl ether lipids as standalone agents and lays the foundation for future studies to elucidate their molecular targets, optimize their design, and evaluate their clinical potential in treating neurodegenerative and inflammatory brain disorders.

## References

1. Braverman, N.E. and Moser, A.B. (2012) Functions of plasmalogen lipids in health and disease. Biochim Biophys Acta 1822 (9), 1442–52.

2. Hossain, M.S. et al. (2017) Reduction of Ether-Type Glycerophospholipids, Plasmalogens, by NF-kappaB Signal Leading to Microglial Activation. J Neurosci 37 (15), 4074–4092.

3. Hossain, M.S. et al. (2020) Biological Functions of Plasmalogens. Adv Exp Med Biol 1299, 171–193.

4. Kim, K.A. et al. (2016) Gut microbiota lipopolysaccharide accelerates inflamm-aging in mice. BMC Microbiol 16, 9.

5. Kumar, A. (2018) Editorial: Neuroinflammation and Cognition. Front Aging Neurosci 10, 413.

6. Benarroch, E. (2024) What Is the Role of Cytokines in Synaptic Transmission? Neurology 103 (8), e209928.

7. Song, B. et al. (2021) Roles of Cytokines in the Temporal Changes of Microglial Membrane Currents and Neuronal Excitability and Synaptic Efficacy in ATP-Induced Cortical Injury Model. Int J Mol Sci 22 (13).

8. Levin, S.G. and Godukhin, O.V. (2017) Modulating Effect of Cytokines on Mechanisms of Synaptic Plasticity in the Brain. Biochemistry (Mosc) 82 (3), 264–274.

9. Khairova, R.A. et al. (2009) A potential role for pro-inflammatory cytokines in regulating synaptic plasticity in major depressive disorder. Int J Neuropsychopharmacol 12 (4), 561–78.

10. Patterson, P.H. and Nawa, H. (1993) Neuronal differentiation factors/cytokines and synaptic plasticity. Cell 72 Suppl, 123–37.

11. Ali, F. et al. (2019) Plasmalogens Inhibit Endocytosis of Toll-like Receptor 4 to Attenuate the Inflammatory Signal in Microglial Cells. Mol Neurobiol 56 (5), 3404–3419.

12. Hossain, M.S. et al. (2022) Plasmalogen-Mediated Activation of GPCR21 Regulates Cytolytic Activity of NK Cells against the Target Cells. J Immunol 209 (2), 310–325.

13. Hossain, M.S. et al. (2022) Plasmalogens, the Vinyl Ether-Linked Glycerophospholipids, Enhance Learning and Memory by Regulating Brain-Derived Neurotrophic Factor. Front Cell Dev Biol 10, 828282.

14. Katafuchi, T. et al. (2012) Effects of plasmalogens on systemic lipopolysaccharide-induced glial activation and beta-amyloid accumulation in adult mice. Ann N Y Acad Sci 1262, 85–92.

15. Petrozziello, T. et al. (2024) Targeting Myeloperoxidase to Reduce Neuroinflammation in X-Linked Dystonia Parkinsonism. CNS Neurosci Ther 30 (11), e70109.

16. Inchiosa, M.A., Jr. (2024) Beta(2)-Adrenergic Suppression of Neuroinflammation in Treatment of Parkinsonism, with Relevance for Neurodegenerative and Neoplastic Disorders. Biomedicines 12 (8).

17. Tong, T. et al. (2023) Corrigendum to “Paraquat exposure induces Parkinsonism by altering lipid profile and evoking neuroinflammation in the midbrain” [Environ. Int. 169 (2022) 107512]. Environ Int 178, 108063.

18. Cunha, D.M.G. et al. (2022) Neuroinflammation in early, late and recovery stages in a progressive parkinsonism model in rats. Front Neurosci 16, 923957.

19. Fujino, T. et al. (2020) Therapeutic Efficacy of Plasmalogens for Alzheimer’s Disease, Mild Cognitive Impairment, and Parkinson’s Disease in Conjunction with a New Hypothesis for the Etiology of Alzheimer’s Disease. Adv Exp Med Biol 1299, 195–212.

20. Fujino, T. et al. (2017) Efficacy and Blood Plasmalogen Changes by Oral Administration of Plasmalogen in Patients with Mild Alzheimer’s Disease and Mild Cognitive Impairment: A Multicenter, Randomized, Double-blind, Placebo-controlled Trial. EBioMedicine 17, 199–205.

21. Hossain, M.S. et al. (2013) Plasmalogens rescue neuronal cell death through an activation of AKT and ERK survival signaling. PLoS One 8 (12), e83508.

22. Hossain, M.S. et al. (2023) Plasmalogens inhibit neuroinflammation and promote cognitive function. Brain Res Bull 192, 56–61.

23. Hossain, M.S. et al. (2018) Oral ingestion of plasmalogens can attenuate the LPS-induced memory loss and microglial activation. Biochem Biophys Res Commun 496 (4), 1033–1039.

24. Sejimo, S. et al. (2018) Scallop-derived plasmalogens attenuate the activation of PKCdelta associated with the brain inflammation. Biochem Biophys Res Commun 503 (2), 837–842.

25. Bligh, E.G. and Dyer, W.J. (1959) A rapid method of total lipid extraction and purification. Can J Biochem Physiol 37 (8), 911–7.

26. Mawatari, S. et al. (2016) Measurement of Ether Phospholipids in Human Plasma with HPLC-ELSD and LC/ESI-MS After Hydrolysis of Plasma with Phospholipase A1. Lipids 51 (8), 997–1006.

27. Mawatari, S. et al. (2012) Dietary plasmalogen increases erythrocyte membrane plasmalogen in rats. Lipids Health Dis 11, 161.

28. Mawatari, S. et al. (2020) Identification of plasmalogens in Bifidobacterium longum, but not in Bifidobacterium animalis. Sci Rep 10 (1), 427.

29. Honsho, M. et al. (2022) ATP8B2-Mediated Asymmetric Distribution of Plasmalogens Regulates Plasmalogen Homeostasis and Plays a Role in Intracellular Signaling. Front Mol Biosci 9, 915457.

30. Hossain, M.S. et al. (2024) KIT-13, a novel plasmalogen derivative, attenuates neuroinflammation and amplifies cognition. Front Cell Dev Biol 12, 1443536.

31. Hossain, M.S. et al. (2016) Neuronal Orphan G-Protein Coupled Receptor Proteins Mediate Plasmalogens-Induced Activation of ERK and Akt Signaling. PLoS One 11 (3), e0150846.

32. Fujino et al. (2018) Effects of Plasmalogen on Patients with Mild Cognitive Impairment: A Randomized, Placebo-Controlled Trial in Japan. J Alzheimers Dis Parkinsonism 2018, 8:1.

33. Fujino et al. (2019) Effects of Plasmalogen on Patients with Moderate-to-Severe Alzheimer’s Disease and Blood Plasmalogen Changes: A Multi-Center, Open-Label Study. J Alzheimers Dis Parkinsonism 2019, 9:4.

